# Volumetric Segmentation and Characterisation of the Paracingulate Sulcus on MRI Scans

**DOI:** 10.1101/859496

**Authors:** Junwei Yang, Duo Wang, Colleen Rollins, Matthew Leming, Pietro Liò, John Suckling, Graham Murray, Jane Garrison, Arnaud Cachia

## Abstract

Many architectures of deep neural networks have been designed to solve specific biomedical problems, among which segmentation is a critical step to detect and locate the boundaries of the target object from an image. In this paper, we develop a deep learning based framework to automatically segment the paracingulate sulcus (PCS) from the MRI scan and estimate lengths for its segments. The study is the first work on segmentation and characterisation of the PCS, and the model achieves a Dice score of over 0.77 on segmentation, which demonstrates its potential for clinical use to assist human annotation. Moreover, the proposed architecture as a solution can be generalised to other problems where the object has similar patterns.

## 1. INTRODUCTION

Over the last decade, deep learning has been widely applied to biomedical researches due to its strong ability of feature extraction. The obtained results can assist analysis on scans to reduce human participation, and can also facilitate treatment.

The PCS is a fold in the cortical surface that lies along the middle of the brain. The sulcus has been proved to be highly related to psychiatric and psychological phenomena such as hallucinations, where 1 cm reduction in sulcal length can increase the chance of hallucination by 20%, semantic fluency, and schizophrenia [1, 2, 3]. The morphology of the sulcus is highly variable between individuals [4], and the its presence can also affect the size of anterior cingulate and paracingulate cortices, which contributes to the control of emotion, inhibitory, and cognition [5, 6]. One of the widely used methods to manually identify the PCS is explained in [1]. Recent researches have demonstrated impressive accuracy for deep neural networks to detect various organs. However, no architectures have been developed and proved to perform well on segmentation of the PCS.

Substantial researches to characterise PCS morphology have been conducted, such as the manual tracing of the sulcus based on certain heuristics [6, 7] using the relative position to the cingulate sulcus, and also the semi-automated scan viewing software (BrainVISA) that can automatically segment the brains sulci with manual intervention [8]. However, both methods require human participation, and the obtained results can be highly dependent on human experiences. Nowadays, deep learning based segmentation architectures have been applied on various types of biomedical images [9, 10]. One of the pioneering models on semantic segmentation task is the fully convolutional neural network, which can predict the object class for each pixel [11]. Inspired by this approach, variants of the network have been developed for biomedical image segmentation. Among them, U-Net is a popular architecture with a U-shaped structure and consists of two paths, namely the contraction and the expansive path [12]. Based on the U-Net, further improvements have also been achieved by adding short and long skip connections [13], and the structure is also altered to allow segmentation on 3D images [14]. More architectures have been proposed based on the fully convolutional network to accelerate convergence. The Deep Contour-Aware Network (DCAN), introduced the concept of intermediate contour label to improve convergence during training [15]. Later, the Coarse-to-Fine Stacked Fully Convolutional Network was developed to segment lymph nodes in the ultrasound images, with a coarse-to-fine pipeline [16].

Existing models are good at finding objects with distinctive visual features, however, the PCS is morphologically similar to other grooves in the brain and only appears in a specific region. As a result, advanced approaches are required to encode its context. In order to overcome the difficulty, a multistage model is proposed in this paper. A coordinate detector is firstly applied to locate the region where the PCS is most likely to appear, then the segmentation network can focus on that region to make predictions. With the predicted segmentation mask, an approach inspired by the shortest path algorithm is used to identify PCS segments and produce length estimates for segments in each brain hemisphere.

In this paper, we develop an end-to-end framework to generate desired information upon presence of the PCS, introduce an error metric to measure the discrepancy between the actual and estimated length, and test its performance across various hyper-parameters. The tool is easy to use, and the code is available upon request from the corresponding author.

## 2. METHODS

### 2.1. Architectures

There are three components in the architecture, the coordinate detector that can find the centre of areas that contain the PCS, the segmentation network that can identify voxels of PCS segments, and the length estimator that can approximate the length of the PCS for each brain hemisphere.

The coordinate detector is a simple convolutional neural network that predicts the centre coordinate of all PCS segments. The network has two convolutional layers with the kernel size of 3 followed by dropout and fully connected layer. Then the image can be cropped around the predicted centre with a pre-determined box size and used as the input to segmentation network, which can not only improve the efficiency of the algorithm but also avoid incorrect identification.

The neural network model used for segmentation is a 3D variant of the U-Net [14]. More specifically, the input only has one channel as the MRI scan is provided in greyscale, and there are two channels in the output indicating whether the voxel correspond to the PCS.

The length estimator provides mathematical based methods to approximate the length of the PCS. First, given the segmentation mask, all isolated groups of PCS voxels are identified. Next, the furthest two points in the 3D space are located for each segment of the complete line. Then, the distance between two points is calculated as estimation. Lastly, the obtained distances in all groups are summed and reported as the length of the PCS for left and right brain hemisphere.

### 2.2. Dataset

The dataset is provided by the Brain Mapping Unit^1^, including 307 T1 structural MRI scans with no identifying information from a group of patients, and 614 manually annotated PCS labels in NIFTI format with the size of 182 × 218 × 182.

The original dataset is used to train the coordinate detector, and images are cropped to train the segmentation network. However, the sample size is not large enough for training a complex model. To prevent from overfitting, we used the data augmentation methods of adding small random perturbations to the cropping coordinates. Translation in seven directions by distance ranged from 0 to 25 with an interval of 1 voxel is applied to generate 10,606 valid images.

The length for segments in each MRI scan is generated based on the tracings on the original T1 scans. The actual centre coordinates of all PCS segments are also computed as their geometric centre.

We use a 8:1:1 ratio to split datasets into training, validation and test set. We use validation dataset to grid search for the best model and test dataset to report evaluation results.

### 2.3. Training

#### 2.3.1. Sliding Window

Considering that the PCS only occupies only a small proportion of the image, a sliding window is applied to crop the scan into a patch that contains segments of the PCS. In the experiment, the window size is chosen to be able to just cover the largest segment in the dataset, which is 64 × 77 × 93.

#### 2.3.2. Loss Function

When training of the coordinate detector, the mean square error is used as the loss function, which is:

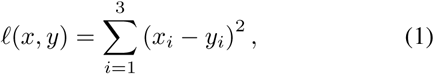

where *x* and *y* are two 3D coordinates, and can be interpreted as the square of the distance between the actual and the predicted centre coordinate.

As for the segmentation network, the weighted binary cross entropy loss is applied. The binary cross entropy loss is a widely used loss function based on the concept of entropy, representing the higher similarity between two distributions.

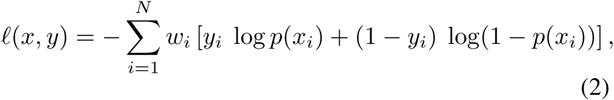

where *w*_*i*_ is the class weight for the *i*-th voxel, in the experiment a 85:15 weight ratio is used to prioritise the PCS voxels.

#### 2.3.3. Metrics

The Dice coefficient averaged across all channels is applied to evaluate the segmentation network, and is defined as:

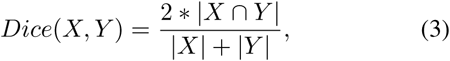

which measures the similarity between two sets *X* and *Y*. If set *X* is identical to set *Y*, the coefficient equals to 1, whereas it equals to 0 when set *X* and set *Y* has no common elements. In other cases, the score will be between 0 and 1.

On the other hand, the absolute percent error is used to measure the accuracy of length estimation. The error is reported as a percentage averaged across all images. Suppose the actual and the estimated length is *l*_*a*_ and *l*_*e*_ respectively, the error can be computed as:

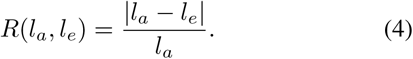

### 2.4. Platform

The package used for training is PyTorch. During training, the Adam optimiser is chosen to update model weights with the initial learning rate of 0.0002 for both networks. The coordinate detector and the segmentation network are trained for 20 and 10 epochs respectively. Due to the large size of the 3D image, a batch size of 2 is used to ensure that the allocated hardware resources are enough to load each batch.

## 3. RESULTS

### 3.1. Coordinate Detector

The coordinate detector is directly trained on a subset of unmodified dataset where images without the PCS are discarded, and evaluated on the images from the test set.

The average mean square error obtained on the test set is 363.7, which is about 19 voxels. An illustration of the prediction is demonstrated in Figure 2. In the figure, the small region that contains the PCS is enlarged for easier visualisation, and the predicted position differs slightly from the actual centre by 10 voxels. Sometimes the predicted position may be further from the centre, a relatively large sliding window size can ensure most of the PCS is covered within the area.

**Fig. 1.**
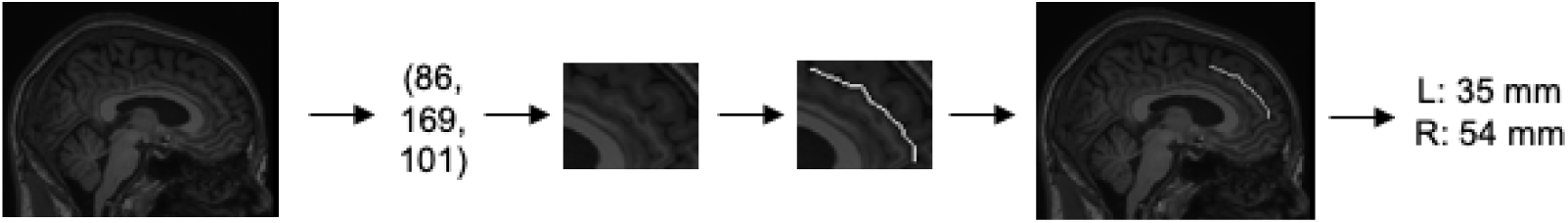
The overview of the framework, in which images and exact numbers are for illustrative purposes only. The centre of the area that may contain the PCS is firstly produced as a coordinate, then the image is cropped around the centre and segmented to mark PCS segments. Based on the prediction, lengths of segments in both brain hemispheres are estimated.

**Fig. 2.**
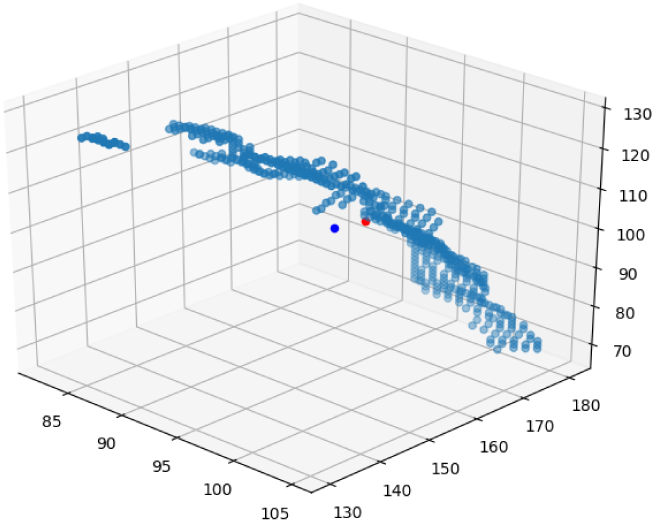
An example of the prediction made by the coordinate detector. Two axes correspond to the *y* and *z* axis, and the *x* axis is fixed. Dots in light blue represent the PCS voxels, and the blue and red dots are the predicted centre and the real centre respectively.

### 3.2. Segmentation Network

With the position predicted by the coordinate detector, the window around the centre can be cropped and fed into the segmentation network. To evaluate the effectiveness of the coordinate detector, images cropped around the fixed average centre position across all segments and around the true position for each segment are also used to train the model.

As shown in Table 1, model using the coordinate detector slightly outperforms the results using the average centre coordinate, and predictions with the correct coordinate have the highest score, which demonstrates the potential of the coordinate detector.

**Table 1.** The Dice score obtained after training the segmentation network on the dataset cropped around different centres.

| Type | Dice score % |
| --- | --- |
| Fixed coordinate | 77.34 |
| Use of coordinate detector | 77.43 |
| Use of true coordinate | 78.10 |

To compare with direct use of state-of-the-art models, training is also performed on the whole image to evaluate the mechanism of sliding window. It turns out that the obtained Dice score is much lower and the model failed to capture the morphological information of the PCS. An example of a slice predicted by the model is displayed in Figure 3. It can be seen that the model tries to label grooves in different parts of the brain as the PCS, and some of the predictions are discontinuous and do not follow the pattern mentioned in the measurement protocol.

**Fig. 3.**
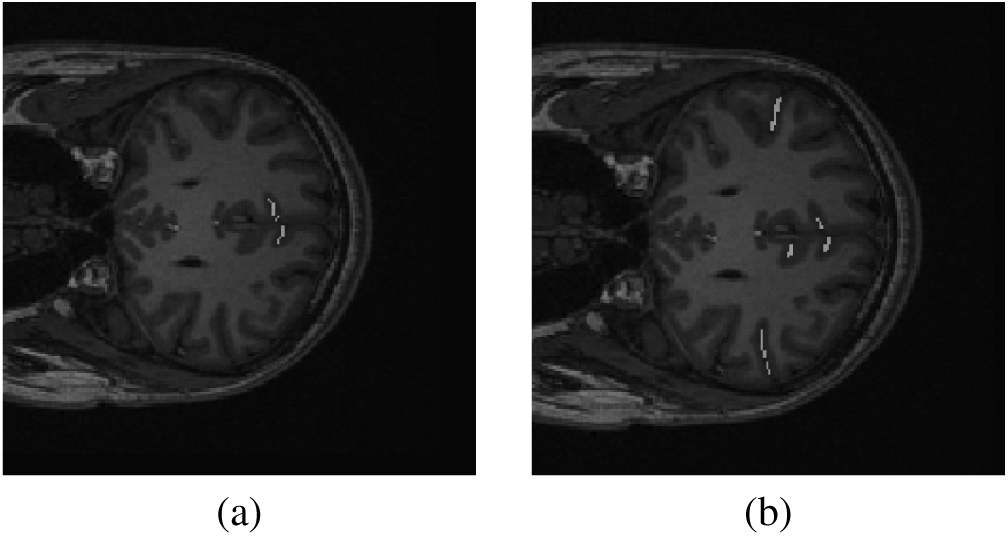
An example of the prediction made by the model trained on whole images. (a) ground truth; (b) prediction.

An example with the Dice score of 82.57% is shown in Figure 4, in which the predicted label is quite accurate and can take significantly less time to annotate the images than a human expert.

**Fig. 4.**
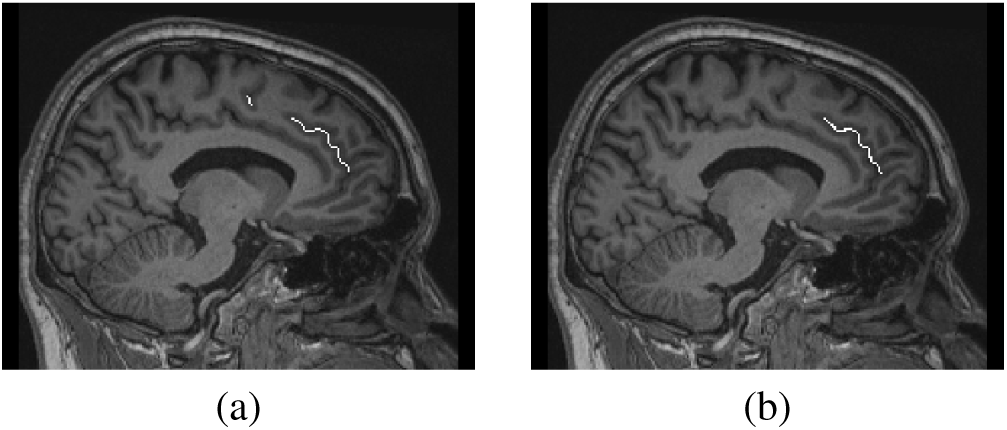
An example slice of the prediction made on a randomly selected MRI scan. (a) ground truth; (b) prediction.

To explore other possible architectures, certain parts of the network are altered to search for optimal structure. Firstly, the kernel size of each convolutional layer being 1 and 5 is attempted for comparison so that the model can learn to extract high-level features at different levels. A smaller kernel allows the model to pay attention to local details of the image, while a larger kernel can capture more general features of the image. Another type of exploration on the architecture is to replace the two consecutive convolutional layers with the residual block used in [17].

The score on the test set is listed in Table 2. As shown in the table, compared to the original model, the score drops dramatically when using the kernel with the size of 1, indicating that features considering contents in surrounding voxels are necessary to identify the PCS due to its morphological pattern.

**Table 2.** The Dice score of the modified model on the test set. The modification types include changes in the kernel size and the replacement of residual block.

| Type | Dice score % |
| --- | --- |
| Kernel (1) | 51.57 |
| Kernel (3) | 77.43 |
| Kernel (5) | 76.57 |
| Residual block | 74.12 |

As for the model with a larger kernel, it turns out that the model with a kernel size of 5 has a slightly lower Dice score, which shows that the kernel with a medium size has a better balance between focusing on details of the PCS and learning the pattern compared to its neighbouring brain tissues.

Skip connections are used to reconstruct the information from the initial layers and can achieve faster traverse of the information in networks. The result shows that the model is outperformed by the original model with a kernel size of 3 and 5. The power of residual connections normally start to appear when the network becomes very deep, and architecture with such connections is effective in capturing features that require learning long-term dependencies.

### 3.3. Length Estimator

The last step of the framework is to produce the estimated length based on the prediction of segments. To evaluate its performance, the average error is computed between estimated and provided lengths, and algorithm is applied to ground truths and predictions of segments.

The results are demonstrated in Table 3. The average length across all segments are 41.70 mm, and 56.06 mm across images in the test set. In the table, the errors are quite consistent between left and right hemispheres at around 17%, and for ground truths from the test set, the average error is quite close to the that evaluated on all images.

**Table 3.** The error obtained by running the estimation algorithm on whole scan images from the dataset, where LH and RH stand for left hemisphere and right hemisphere respectively.

| Dataset | Overall % | LH % | RH % |
| --- | --- | --- | --- |
| Ground truth (all) | 16.87 | 16.84 | 16.89 |
| Ground truth (test set) | 16.41 | 16.99 | 15.90 |
| Prediction (test set) | 21.63 | 22.67 | 20.60 |

The performance is quite competitive compared to the error acquired on ground truths from the test set. After closer investigation of the generated masks and consultation with experts, the prediction is generally accurate, and the algorithm can successfully produce the length for each segment. However, there are cases where PCS segments cannot be found by the segmentation network, which leads to 100% error for that sample, and can increase the average error significantly.

## 4. CONCLUSION

In this paper, we develop an automated tool to identify and characterise the PCS. The framework considers the particular morphology of the PCS and utilises the CNN-based model to perform segmentation on the region of interest. The configuration used for training and several mechanisms is specifically designed to improve the performance compared to direct use of other state-of-the-art models. The experimental results demonstrate that the model performs well upon the presence of the PCS, at an acceptable level of the error, and most importantly, provides an approach that can be applied to morphologically similar objects.

## 5. ACKNOWLEDGEMENT

We would like to acknowledge the help from research students who prepared the dataset used in this paper, including Aida Seyed-Salihi, Maite Arribas, Aniket Patel, Rory Durham, and Charlotte Jones. This work was supported by Gates Cambridge and the H.E. Durham Fund.

## Footnotes

1 Brain Mapping Unit, Department of Psychiatry, University of Cambridge.

